# The effective multiplicity of infection for HCMV depends on activity of the cellular 20S proteasome

**DOI:** 10.1101/2024.10.21.619499

**Authors:** Katie M. Cataldo, Kathryn L. Roche, Christopher E. Monti, Ranjan K. Dash, Eain A. Murphy, Scott S. Terhune

## Abstract

Human cytomegalovirus (HCMV) is a betaherpesvirus capable of infecting numerous cell types and persisting throughout an infected individual’s life. Disease usually occurs in individuals with compromised or underdeveloped immune systems. Several antivirals exist but have limitations relating to toxicity and resistance. HCMV replication involves upregulation of host proteasomal activities which play important roles in the temporal stages of replication. Here, we defined the impact on replication kinetics of the proteasome inhibitor, bortezomib. We demonstrate that bortezomib significantly reduces levels of viral genomes and infectious virions produced from a population of cells. Inhibition reduced expression of viral proteins that are influenced by genome synthesis. When added prior to 24 hpi, we observe decreases in PCNA and Cdk1 while increases in p21 whose regulations contribute to efficient replication. This response synergized with an antiviral, maribavir. Since some replication occurred, we tested the hypothesis that a subset of infected cells might break through inhibition. Initially, we simulated bortezomib activities using a mechanistic computational model of late-lytic replication. Upon reducing MOI *in-silico*, we observed near identical simulated results compared to experimental data. Next, we analyzed replication using live-cell imaging. This revealed treated cultures do contain a population of cells with fully developed late-stage cytoplasmic assembly compartments but at significantly lower numbers. We refer to this as the effective MOI. Overall, our studies support a hypothesis in which 20S proteasome inhibition disrupts HCMV replication by reducing the MOI to an effective MOI, defined by a fraction of infected cells capable of progressing to fulminant infection.

**IMPORTANCE:** HCMV infection and reactivation continues to contribute to morbidity and mortality around the world. Antiviral compounds are available but have limitations. Here, we have defined the impact of the proteasome inhibitor bortezomib on HCMV replication. Proteasomal activities play a critical role in temporal changes required for replication. We demonstrate that disrupting these activities inhibits viral replication while likely supporting increased antiviral activity of the anti-HCMV agent, maribavir. Using a combination of live cell imaging and computational tools, we discover that a subset of infected cells progresses to fulminant infection which we define as the effective MOI, and this subset would otherwise be missed when analyzing the average of the population.

## INTRODUCTION

Human cytomegalovirus (HCMV) is in the *betaherpesvirinae* subfamily of herpesviruses which also include HHV-6A, -6B and -7 (Reviewed in (1)). HCMV infects numerous cell types and tissues throughout the human body. Following an initial exposure, infection becomes life-long consisting of a balance between lytic replication and latent maintenance states. Life-threatening disease can occur upon reduced or absence of host immunity frequently observed in solid organ and bone marrow transplants. There are currently no vaccines available against HCMV which continues to be a priority area in vaccine development (2). Several antiviral compounds are approved to manage HCMV infection targeting different steps in viral replication (Reviewed in (3)). These includes ganciclovir (a guanine analogue), cidofovir (a cytosine analogue) and letermovir (a non-nucleoside 3,4-dihydroquinazolinyl acetic acid). Ganciclovir, and its oral equivalent valganciclovir, is a nucleoside analogue phosphorylated by the HCMV pUL97 kinase resulting in ganciclovir-5-triphosphate which has a high affinity for incorporation by the viral polymerase thereby terminating genome synthesis. Similarly, cidofovir is phosphorylated by a cellular kinase also resulting in termination of genome synthesis upon incorporation. Letermovir inhibits replication by associating with the pUL56 subunit of the viral terminase complex and disrupting encapsidation and cleavage of concatemeric genomes. An additional compound being used is the antiviral maribavir, a benzimidazole riboside which is a selective ATP competitor of the pUL97 kinase disrupting activities primarily at virion egress.

HCMV infection involves a protracted lytic replication period requiring temporal changes in viral and host gene expression as well as cellular processes and compartments. Following entry, the viral tegument proteins are deposited within the cytoplasm and the capsid translocates to the nucleus where the genome is deposited. Initiation of gene expression progresses in a coordinated process involving expression of immediate early (IE), early (E), early-late (E-L), and true late (L) RNAs, with late expression dependent on viral DNA synthesis. Similarly, temporal patterns of viral protein levels have been defined by Weekes et al. (4) which include Tp1 (temporal profile), Tp2, Tp3/4, and Tp5. These RNAs and proteins participate in altering cellular processes and compartments required to support virion production. HCMV proteins disrupt cellular processes such as intrinsic immune responses, cell cycle, host gene expression, and cellular metabolism. These altered cellular network interactions are accompanied by changes in cellular compartments and protein composition (5). Prominent newly formed intracellular compartments include a nuclear replication compartment which is a membrane-less region defined by newly synthesized viral genomes, and a late stage cytoplasmic assembly compartment (6).

The temporal changes in protein levels and interaction networks are dependent on host cellular proteasome activities (Reviewed in (7)). The 26S proteasome is responsible for most of the ubiquitin-dependent protein degradation and consists of a 20S catalytic core complex and one or two 19S regulatory complexes. In contrast, 20S ubiquitin-independent activity is regulated by the 11S regulatory complex. The 20S catalytic core is a multi-subunit structure with caspase-like, trypsin-like, and chymotrypsin-like activities. Inhibitors of 20S activities are now available for clinical use in cancer therapies including the reversible inhibitor, bortezomib which targets the chymotrypsin-like activity. During the progression of HCMV infection, proteasome activity and subunit expression levels increase (8) which play an important role in modulating levels of viral protein classes (9). In addition, numerous HCMV proteins function by inducing host protein degradation in a proteasome-dependent manner. Examples include HCMV pp71-stimulated degradations of Rb, Daxx, and BclAF1 (10–12), pUL27-targeted Tip60 (13, 14) and pUL21-mediated disruptions of APC/C and cyclin A (15, 16). Infection induces degradation of additional host restriction factors involving both proteasome-dependent and independent mechanisms (9). Upon the addition of pan-proteasome inhibitors, HCMV replication is disrupted at the level of viral gene expression and genome synthesis which also occurs when disrupting expression of specific proteasome subunits (8). Interestingly, the 19S regulatory complex has been implicated in stimulating viral gene expression independent of the catalytic core activity (8, 17).

In this study, we demonstrate that the 20S proteasome inhibitor bortezomib disrupts viral genome synthesis, impacting viral gene expression and virion production as well as levels of key cell cycle factors. We provide evidence that bortezomib likely synergizes with the antiviral compound, maribavir to block infection which is based on a previous hypothesis (14) relating p21 levels to antiviral efficacy. Finally, by combining in silico analysis using a computational model of late lytic replication with live cell imaging, we reveal that a fraction of cells progresses to fulminant infection.

## RESULTS

### Bortezomib disrupts HCMV viral DNA synthesis and early and late protein expressions impacting viral titers

The host proteasome plays an important role in the protracted life cycle of HCMV. Therefore, we were interested in testing the impact of the proteasome inhibitor, bortezomib on HCMV replication. The dipeptide compound bortezomib (BTZ) reversibly inhibits the chymotrypsin-like catalytic activity of the 20S complex, associating with PSMB5 subunit and suppressing proliferation of numerous cancer types. To identify a maximum *in vitro* tolerable concentration, we measured the percent viable and total number of MRC-5 fibroblasts over a 96 hr period using trypan blue exclusion. We treated subconfluent MRC-5 cells with 10, 15, or 25 nM bortezomib or DMSO vehicle control which were replaced every 24 hr with new media and compound (**Fig. 1A**). The highest concentration of 25 nM resulted in a 35% reduction in cell viability by 96 hr and loss of cell division. At 15 nM of bortezomib, we observed minimally reduced proliferation with a small reduction in cell viability (**Fig. 1A**). We selected 15 nM for the remainder of our studies. In patients receiving bortezomib, this is below the initial maximal concentration of 218 nM (83.9 ng/ml) but above the sustained concentration of approximately 2 nM (1 ng/ml) observed over several days (18).

**Figure 1.**
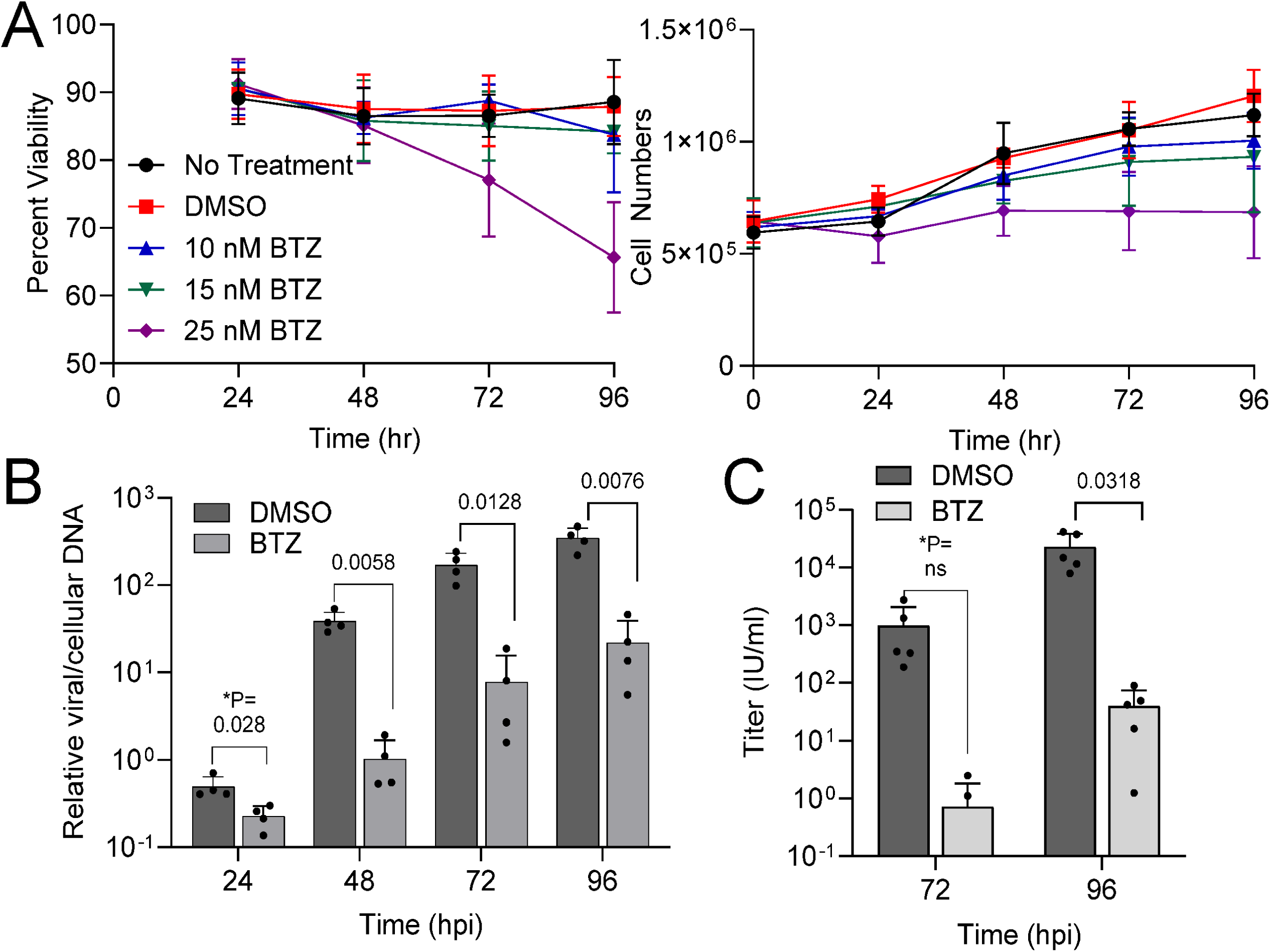
The 20S proteasome inhibitor bortezomib disrupts HCMV genome synthesis and extracellular virus production. **(A)** Subconfluent human MRC-5 lung fibroblasts were treated using the indicated concentrations of bortezomib or DMSO vehicle control. Percent viable cells and total viable cells were determined by trypan blue stain exclusion. Error bars represent standard deviations (n=3). The remainder of the studies used 15 nM bortezomib in DMSO unless otherwise noted. **(B)** Fibroblasts were grown to confluency then infected using HCMV strain TB40/E expressing eGFP (TB40/E-eGFP) at an MOI of 0.5 IU/cell. Samples were treated with 15 nM bortezomib or DMSO control at 2 hpi and changed every 24 hrs afterwards. Culture media and cell pellets were collected at indicated time points. Viral genomes were quantified from whole cell DNA using qPCR and primers to the viral gene UL123 and cellular gene β-tubulin (n=4). Error bars represent standard deviations. **(C)** Culture media were used to determine viral titers by plating serial dilutions, staining for IE1, and counting IE1-postive cells per well at a given volume (n=5). *P values were calculated using a paired T test between control and treated samples for each time point.

To evaluate the effect of bortezomib on replication, we infected confluent fibroblasts here and in the remainder of experiments at an average multiplicity of infection (MOI) of 0.5 infectious units per cell (IU/cell) using TB40-BAC4-derived HCMV expressing eGFP (19, 20). Infection exhibits a Poisson distribution in which some cells may become infected with multiple IU at the expense of other cells. We added 15 nM bortezomib or DMSO vehicle control starting at 2 hpi and changed media and compound every 24 hr. We measured cell-free viral titers and cell-associated viral DNA levels from the population of infected cells at the indicated time points. The addition of bortezomib resulted in reduced viral DNA by 24 hpi with an average 1.5-log reduction by 48 hpi (**Fig. 1B**). Viral DNA levels did increase under both conditions out to 96 hpi yet bortezomib-treated infections remained significantly lower than control. We measured extracellular viral titers which showed a 3-log reduction compared to vehicle at 72 and 96 hpi (**Fig. 1C**). These data indicate that addition of the 20S proteasome inhibitor bortezomib to a population of infected cells significantly reduces levels of viral genomes and production of cell-free infectious virion during lytic replication.

To advance our understanding of the disruption, we analyzed changes in viral protein steady-state levels. We infected confluent fibroblasts as described earlier and analyzed expression levels of HCMV IE1, IE2, pUL44 and pp28 proteins using immunoblot analyses of whole cell lysates from the culture dish collected at multiple time points (**Fig. 2A**). We quantified antibody signal intensities of individual proteins normalized to total protein, and we compared 15 nM bortezomib-treated samples to vehicle-treated control. The immediate early gene product, IE1p72 exhibited similar levels out to 72 hpi with a slight drop at 96 hpi (**Fig. 2B**). We quantified IE1 RNA levels by qPCR and primers to exon 4 which showed decreasing RNA levels reaching significance by 48 hpi and afterwards (**Fig. 2B**). In contrast to IE1, we observed significant reductions in IE2p86 and p60 proteins and IE2 exon 5-containing RNAs by 48 hpi and afterwards in the population (**Fig. 2C**) which is consistent with greater influence of vDNA synthesis on IE2 levels. We also detected reductions in the early-late gene product, pUL44 and the true-late gene product, pp28 starting at 24 hpi and afterwards (**Fig. 2D**). In general, maximal expressions of IE2, pUL44, and pp28 are dependent on HCMV genome synthesis. Further, IE2 and pUL44 are both necessary for genome synthesis while pp28 is necessary for virion assembly (Reviewed in (21)). Overall, our data demonstrate that specific inhibition of the 20S proteasome starting at 2 hpi disrupts replication with observed changes consistent with an overall inhibition of viral genome synthesis in the culture. Our results are consistent with published studies on MG132 inhibition of proteasome activity and HCMV replication (8, 22).

**Figure 2.**
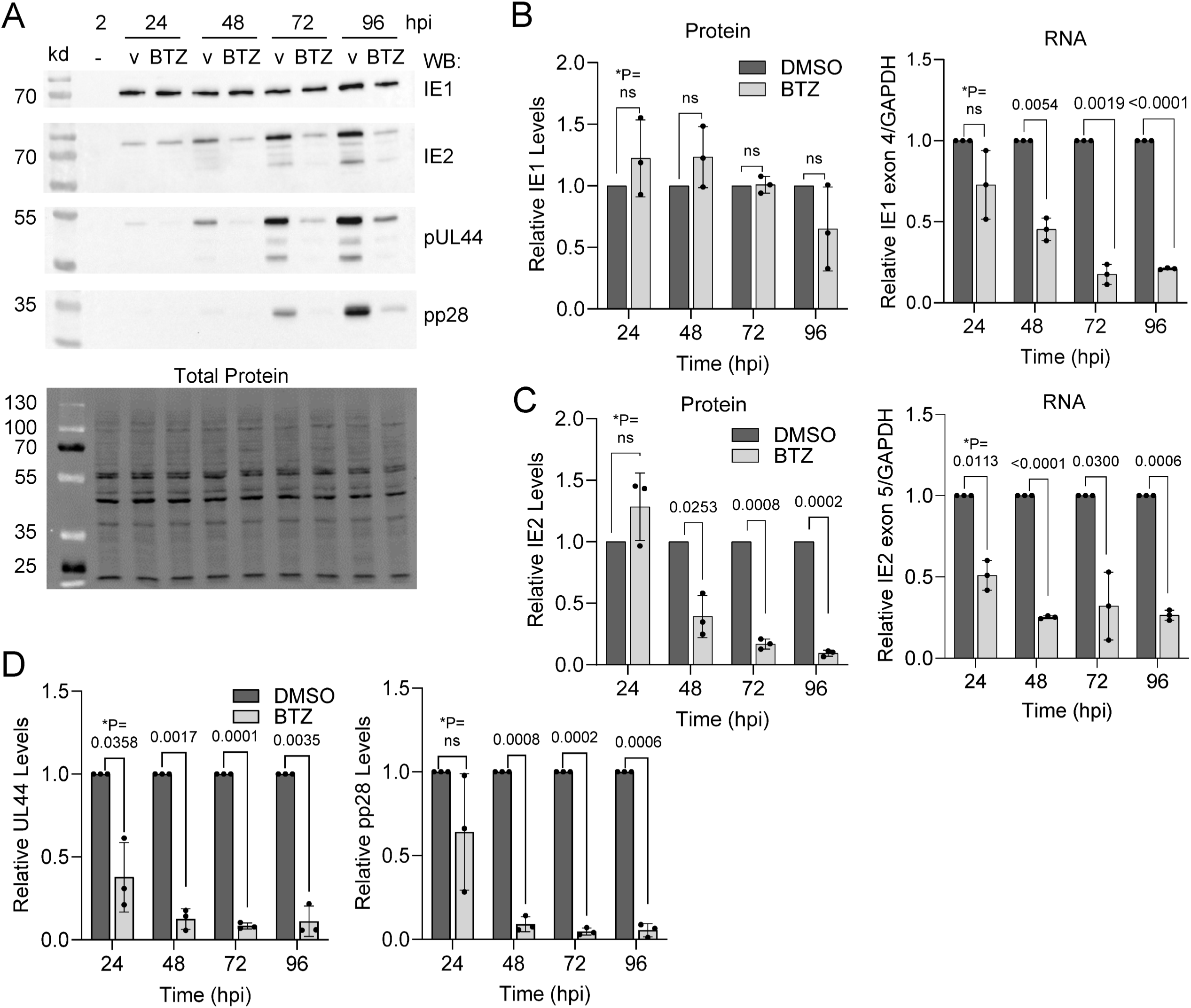
Activity of the 20S proteasome is necessary for HCMV early and late gene expression. **(A)** MRC-5 fibroblasts were grown to confluency and infected at an MOI of 0.5 IU/cell with HCMV TB40/E-eGFP. Starting at 2 hpi, 15 nM bortezomib or DMSO vehicle was added and changed every 24 hrs. Protein expression was analyzed by immunoblot using whole cell lysates isolated at a given time and the indicated antibodies with representative images shown. Total protein levels were detected for each immunoblot using stain-free imaging. **(B**-**D)** Expression levels were determined for **(B)** IE1 protein and RNA, **(C)** IE2 protein and RNA, and **(D)** pUL44 and pp28 protein normalized to total protein per lane. Quantification of protein signal was done using all visible bands in (A), and protein and RNA data are presented as bortezomib-treated relative to vehicle control for each time point (n=3). Error bars represent standard deviation. *P values were calculated using a paired T test between control and treated samples for each time point.

### Early addition of bortezomib disrupts virus-induced changes in cell cycle factors

HCMV replication requires progression through unique cell cycle phases regulated by expressions of numerous viral proteins (Reviewed in (23)) and culminating in a pseudo-mitotic state (24). Further, proteasome activity is an essential activity in regulating ubiquitin-dependent cell cycle oscillations (e.g., APC/C, SCF, etc.). Next, we analyzed the impact of bortezomib on HCMV dysregulation of cell cycle proteins. Confluent MRC-5 fibroblasts were infected at an MOI of 0.5 and treated using 15 nM bortezomib or vehicle control starting at 2 hpi. We limited our analysis to markers of late stage infection including PCNA which accumulates within nuclear replication compartments (25), and Cyclin B-Cdk1 which contributes to virion egress (13, 24, 26) (**Fig. 3A**). Addition of bortezomib resulted in significant reductions in the average levels of Cdk1 at all time points. In contrast, we observed a high degree of variability for Cyclin B levels showing some reduction at 48 and 72 hpi (**Fig. 3B**). Similarly, we observed significant reductions in PCNA at all time points (**Fig. 3B**). Progression through infection is accompanied by calpain and proteasome-dependent degradation of p21 (27). Addition of the 20S proteasome inhibitor resulted in significant increases in p21 levels in the culture at all time points (**Fig. 3A,C**) which was not observed at the RNA level (**Fig. 3C**). This is consistent with the observation that bortezomib stabilizes p21 and p27 in tumor lines (28). During infection, Cyclin B-Cdk1 contributes to capsid egress by disrupting nuclear lamina (13, 26), and we previously hypothesized that increased p21 and reduced Cdk1 levels might alter levels of phosphorylated lamin A/C. CyclinB:CDK1-mediated and pUL97-mediated phosphorylation of lamin A/C induce lamin A/C depolymerization and nuclear lamina breakdown during mitosis or pseudo-mitosis, respectively (13, 26). Therefore, we measured total lamin A/C and phosphorylated lamin A/C at serine 22. However, we did not observe any significant changes (**Fig. 3A,D**) which is contrary to our original hypothesis and suggests a more generalized role of p21 in inhibiting replication. Overall, we conclude that bortezomib disrupts HCMV-mediated increases in levels of PCNA and Cdk1 while preventing HCMV-mediated decrease in cell cycle inhibitor p21 in the population.

**Figure 3.**
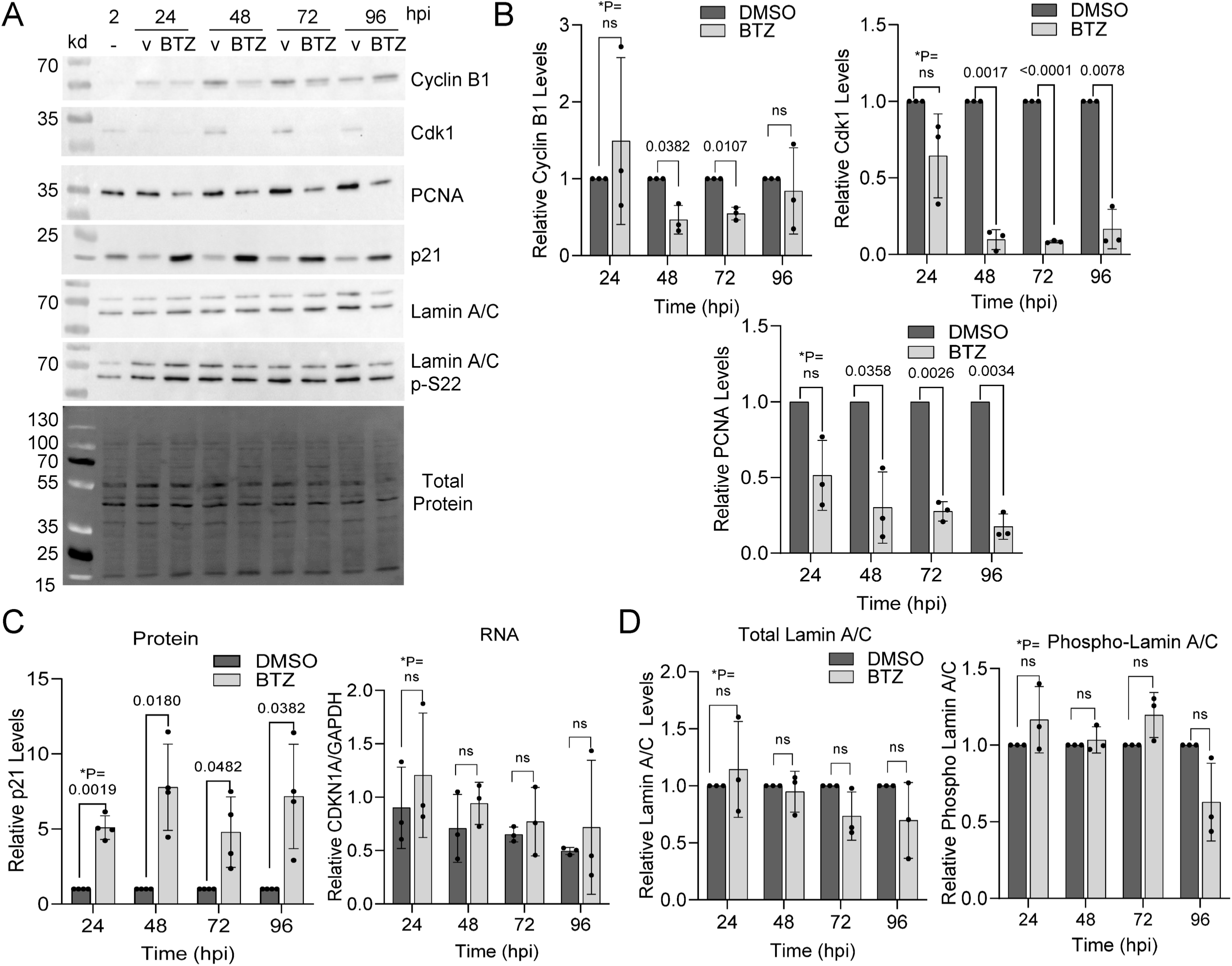
Proteasome inhibition disrupts infection-induced changes to cell cycle regulators. **(A)** Fibroblasts were grown to confluency and infected using 0.5 IU/cell of HCMV TB40/E-eGFP. At 2 hpi, 15 nM bortezomib or DMSO vehicle was added and changed every 24 hrs. Protein expression was analyzed by immunoblot using whole cell lysates and the indicated antibodies. Total protein levels were detected using stain-free imaging and representative images are shown. Expression levels of **(B)** Cyclin B1, CDK1 and PCNA were measured at the indicated times, normalized to total protein per lane, and presented as bortezomib-treated samples relative to vehicle control for each time point (n=3). Error bars represent standard deviation. **(C)** Expression of p21 RNA and protein levels were measured at the indicated times. Protein levels were obtained by normalizing p21 to total protein levels (n=4). RNA levels were determined using RT-qPCR and primers to the p21 gene and normalized to GAPDH (n=4). **(D)** Expression was determined for both bands of Lamin A/C or phosphorylated Lamin A/C at serine 22 levels normalized to total protein of treated relative to untreated conditions are shown (n=3). *P values were calculated using a paired T test between control and treated samples for each time point.

We next asked whether the timing of inhibition would influence outcomes by staggering the addition of 15 nM bortezomib between 2 and 72 hpi, then analyzing culture samples at an endpoint of 96 hpi (**Fig. 4A**). When bortezomib was added at 2 and 24 hpi, we again observed significant decreases in both viral genomes and titers measured at 96 hpi (**Fig. 4B**). Decreases in vDNA were observed when added at 48 and 72 hpi but not statistically significant while no decrease in resulting titers occurred under those treatment conditions (**Fig. 4B**). As described earlier, we also measured viral and cellular proteins (**Fig. 5A**) using immunoblot analysis. We observed trends of decreased IE2, pUL44 and pp28 levels when adding bortezomib early compared to control (**Fig. 5B**). Likewise, addition of bortezomib early trended toward elevated p21 levels and decreased PCNA (**Fig. 5C**). These data suggest that 20S proteasome activity within the first 24 hr of infection contributes to regulating host and viral protein expressions with several of these factors required for progression of HCMV replication.

**Figure 4.**
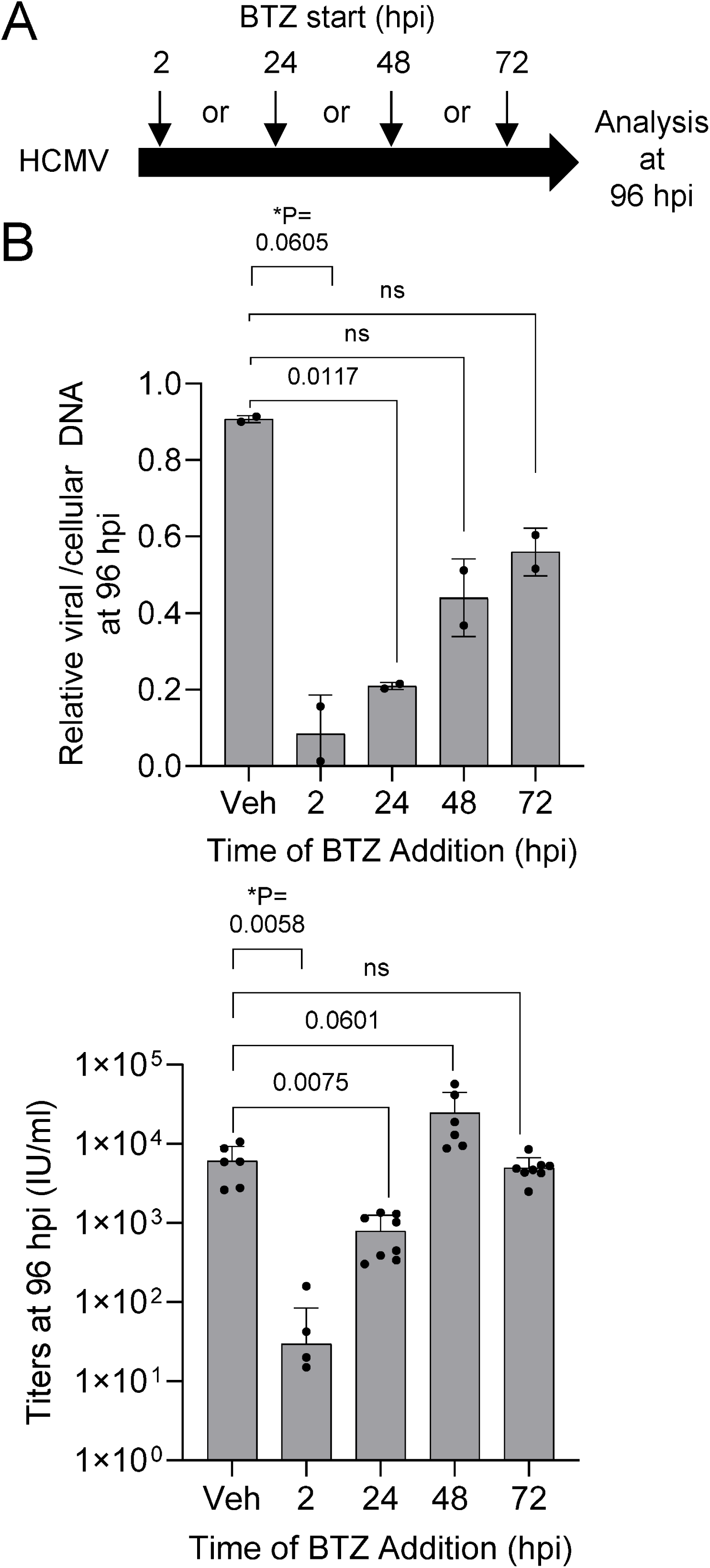
Disruption of HCMV replication depends on the timing of proteasome inhibition. **(A)** Fibroblasts were grown to confluence and infected using 0.5 IU/cell of HCMV TB40/E-eGFP. Cells were washed at 2 hpi and, at the indicated times, 15 nM bortezomib or DMSO vehicle was added and changed every 24 hrs. **(B)** At 96 hpi, culture media were used to determine viral titers by plating serial dilutions, staining for IE1, and counting IE1-postive cells per well at a given volume of inoculum (n=2). HCMV genomes were quantified by qPCR using whole cell DNA and primers to viral UL123 and normalized to cellular β-tubulin (n = 2). Error bars represent standard deviation, and P values were calculated using a paired T test between control and treated samples for each time point.

**Figure 5.**
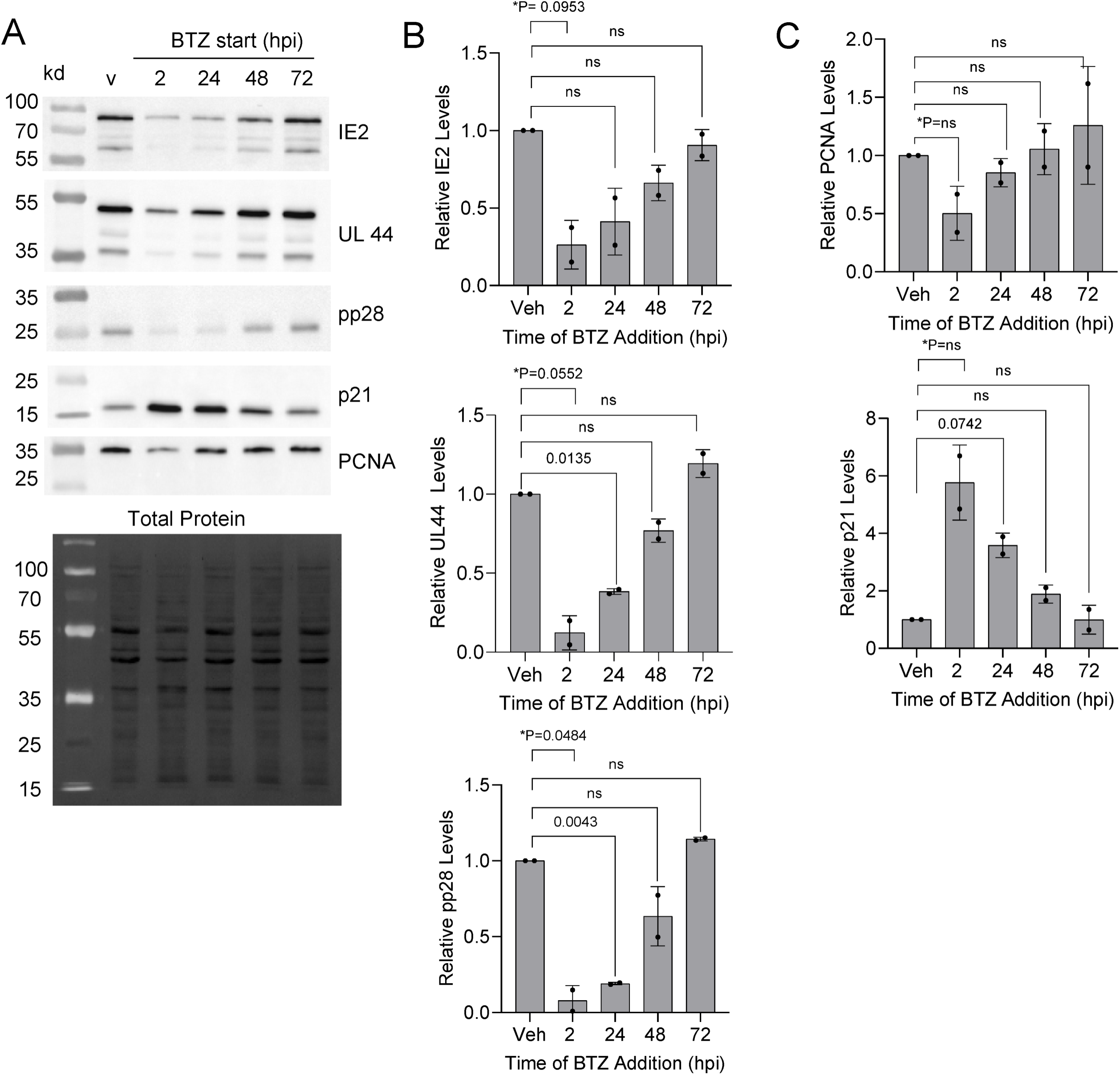
Limited impact to replication when inhibition occurs after 48 hpi. **(A)** Fibroblasts were grown to confluence and infected using 0.5 IU/cell of HCMV TB40/E-eGFP. Cells were washed at 2 hpi and, at the indicated times, 15 nM bortezomib or DMSO vehicle was added and changed every 24 hrs. At 96 hpi, expression levels were determined using immunoblot analysis. **(B,C)** Quantification of protein signal was done using all visible bands, and data are presented as bortezomib-treated relative to vehicle control for each time point for IE2, pUL44 and pp28 and **(C)** PCNA and p21 proteins normalized to total protein per lane (n=2). Error bars represent standard deviation, and P values were calculated using a paired T test between control and treated samples for each time point.

### Bortezomib increases antiviral activity of the HCMV kinase inhibitor maribavir

We have previously demonstrated that compounds causing increased steady-state level of p21 during HCMV infection synergize with the antiviral maribavir (29, 30). Maribavir is a benzimidazole riboside that inhibits the HCMV viral kinase pUL97 (31–33). Having observed a significant increase in p21 (**Fig. 3A,D**), we investigated the potential impact of bortezomib on maribavir antiviral activity. Fibroblast cultures were grown to confluency and infected using 3 IU/cell of HCMV in contrast to 0.5 IU/cell used in earlier studies so that we are establishing an infectivity of at least 99% of the cells in culture. We analyzed varying combinations of bortezomib (0.1, 1.0, 10 nM) and maribavir (0.1, 1.0, 10 µM) on infection. Using 10 nM bortezomib and a multiplicity of 3 IU/cell, we measured an average 0.5-log reduction in viral titers (**Fig. 6A**). Similar reductions were observed using maribavir at 0.1 µM without bortezomib while the addition of 10 µM maribavir resulting in a 2-log reduction (**Fig. 6A**). As we added increasing concentrations of bortezomib to maribavir, we were unable to detect virus in several replicates starting at 0.1 nM bortezomib and 1 µM maribavir (**Fig. 6A**). At 10 nM bortezomib and 10 µM maribavir, only one of the four replicates produced a measurable titer. We also determined changes in viral genomes under these conditions. At a multiplicity of 3 IU/cell, we observed no difference between 10 nM bortezomib and vehicle-treated samples (**Fig. 6B**). We observed an approximate 0.7-log reduction in viral genomes with the addition of maribavir under most conditions. These data demonstrate that combining low level proteasome inhibition with viral kinase inhibition results in a substantial drop in viral titers with limited impact of viral genome levels in the populations.

**Figure 6.**
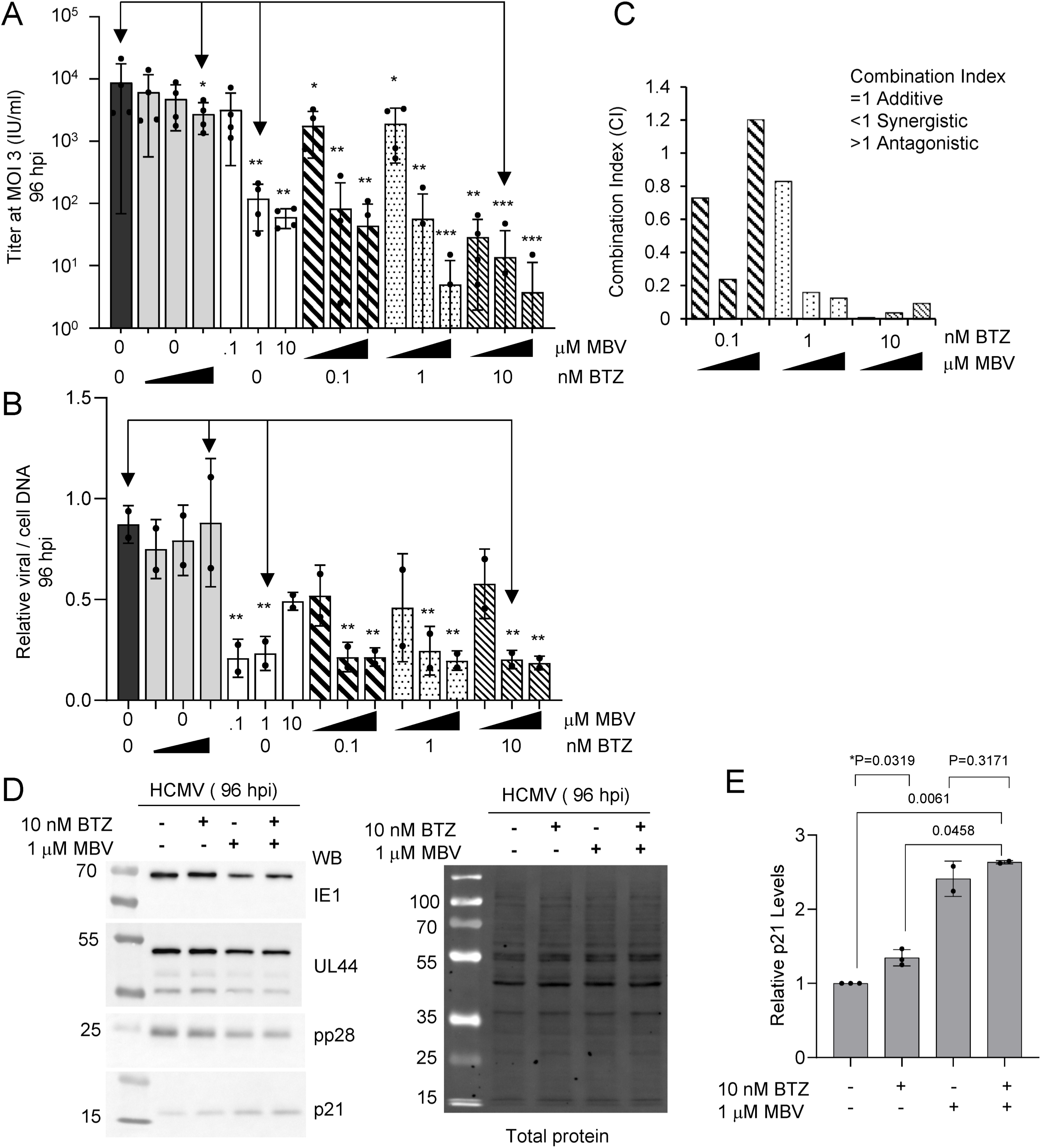
The HCMV kinase inhibitor maribavir behaves synergistically with bortezomib treatment. MRC-5 fibroblasts were grown to confluency and infected using 3 IU/cell of HCMV TB40/E-eGFP. Cells were washed at 2 hpi and varying concentrations of bortezomib (0.1, 1.0 and 10.0 nM), maribavir (0.1, 1.0, 10 µM) or vehicle control were added and changed every 24 hrs. **(A)** Culture media were collected at 96 hpi and viral titers determined by plating serial dilutions, staining for IE1, and quantifying IE1-postive cells per wells at a given volume of inoculum (n=4). Data represent standard deviation and analyzed using one-way ANNOVA with * <0.05. Arrows indicate concentrations tested for protein expressions in (D). **(B)** Whole cell DNA was isolated at 96 hpi and HCMV genomes were quantified by qPCR using whole cell DNA and primers to viral UL123 and normalized to cellular β-tubulin (n=2). Arrows indicate concentrations tested for protein expressions in (D). **(C)** Combination index (CI) values which provide a prediction of how compounds interact were determined using the CompuSyn software tool which utilizes the Chou Talalay Method. These results suggest BTZ and MBV act synergistically at most concentrations. **(D)** Expressions of IE1, pUL44, pp28, and p21 were assessed at 96 hpi using immunoblot analysis following treatment with DMSO, 10 nM bortezomib, 1 µM maribavir, or 10 nM bortezomib and 1 µM maribavir together. Representative images are shown including total protein (n=3). **(E)** Quantification of p21 protein signal normalized to total protein relative to vehicle control for each condition (n=3). Error bars represent standard deviation, and P values were calculated using a paired T test between the indicated samples.

We next asked whether bortezomib and maribavir might act in an additive or synergistic manner to inhibit HCMV. Based on the viral titer data, we calculated Combination Index (CI) values using the software tool, CompuSyn (**Fig. 6C**)(34). Using non-constant ratio combinations of bortezomib and maribavir, the average CI value was determined to be 0.38. However, the values are highly variable. A CI<1 is indicative of synergism, a CI=1 is indicative of an additive effect, and a CI>1 is indicative of antagonism (34). These data suggest, but are not conclusive, that combining low concentrations of bortezomib with maribavir in a cell culture system act synergistically to inhibition HCMV replication.

Finally, we asked whether viral and cellular protein expressions are differentially altered between 10 nM bortezomib, 1 µM maribavir, or both compounds and these conditions are identified by arrows in **Figures 6A and B**. Confluent cultures of fibroblasts were infected as described above with compounds added at 2 hpi. We analyzed IE1, pUL44, pp28, and cellular p21 levels in the population by immunoblot at 96 hpi (**Fig. 6D**). Except for IE1, notable decreases in viral proteins and increase in p21 occurred (**Fig. 6D**). We quantified signal intensities from the p21 antibody that were normalized to total protein per lane and evaluated relative change between conditions (**Fig. 6E**). This analysis reveals increases in p21 at 1.2-fold for bortezomib, 2.2-fold for maribavir, and 2.6-fold when combined. However, surprisingly, difference in p21 between maribavir only and maribavir with bortezomib at these lower concentrations are not significant (**Fig. 6E**). Overall, our data show that addition of maribavir or bortezomib results in elevated p21 levels and evidence of possible synergistic antiviral activity in an *in vitro* fibroblast culture.

### Proteasome inhibition reduces the number of cells in the population developing pp28-positive replication compartments

We observed reduced viral DNA levels and extracellular titers using sub-cytotoxic levels of proteasome inhibition. These observations raised the question of whether this is the result of a uniformed delay in replication in the population, or whether this is the result of a subset of cells progressing to a fulminant infection. As noted earlier, infection occurs in a Poisson distribution and requires tegument-mediated and proteasome-dependent degradation of intrinsic antiviral factors (e.g., Daxx, PML, BclAF1, etc.)(10–12). Therefore, it is conceivable that a subset of cells will receive greater numbers of viral particles overcoming intrinsic resistance in the presence of bortezomib. To begin testing these possibilities, we simulated infection using a late-lytic computational model of HCMV replication (35). This model was developed and parameterized using experimental data from multiple MOIs (0.1, 0.5, and 5 IU/cell) including protein expressions, viral DNA, and extracellular titers during infection of MRC-5 fibroblasts (**Fig. 7A**). Genome synthesis was empirically determined in the model. Using this computational tool, we simulated bortezomib-mediated inhibition of viral DNA synthesis by reducing the input MOI as this model uses a linear relationship between MOI and input vDNA at 2 hpi (35). We fit simulations of MOI 0.5 (**Fig. 7B**) to our experimental data for pp28 (Tp5_2_) (**Fig. 2D**) and viral titers (**Fig. 1C**). Upon mimicking bortezomib inhibition by reducing the MOI *in silico* by 80%, we observed reduced levels of pp28 and viral titers (**Fig. 7B**), and these simulated data overlapped with our bortezomib experimental data. We now refer to this reduced MOI as the effective MOI (MOI_eff_) with equations accounting for the change added to the previous late-lytic computational model as described in Materials and Methods. Our prior parameterization was completed using a range of MOI, and these simulations support the idea that a subset of cells can complete full lytic replication upon inhibition. However, it is still plausible to obtain similar outcomes by simulating reduced protein synthesis rates within the model at an MOI of 0.5.

**Figure 7.**
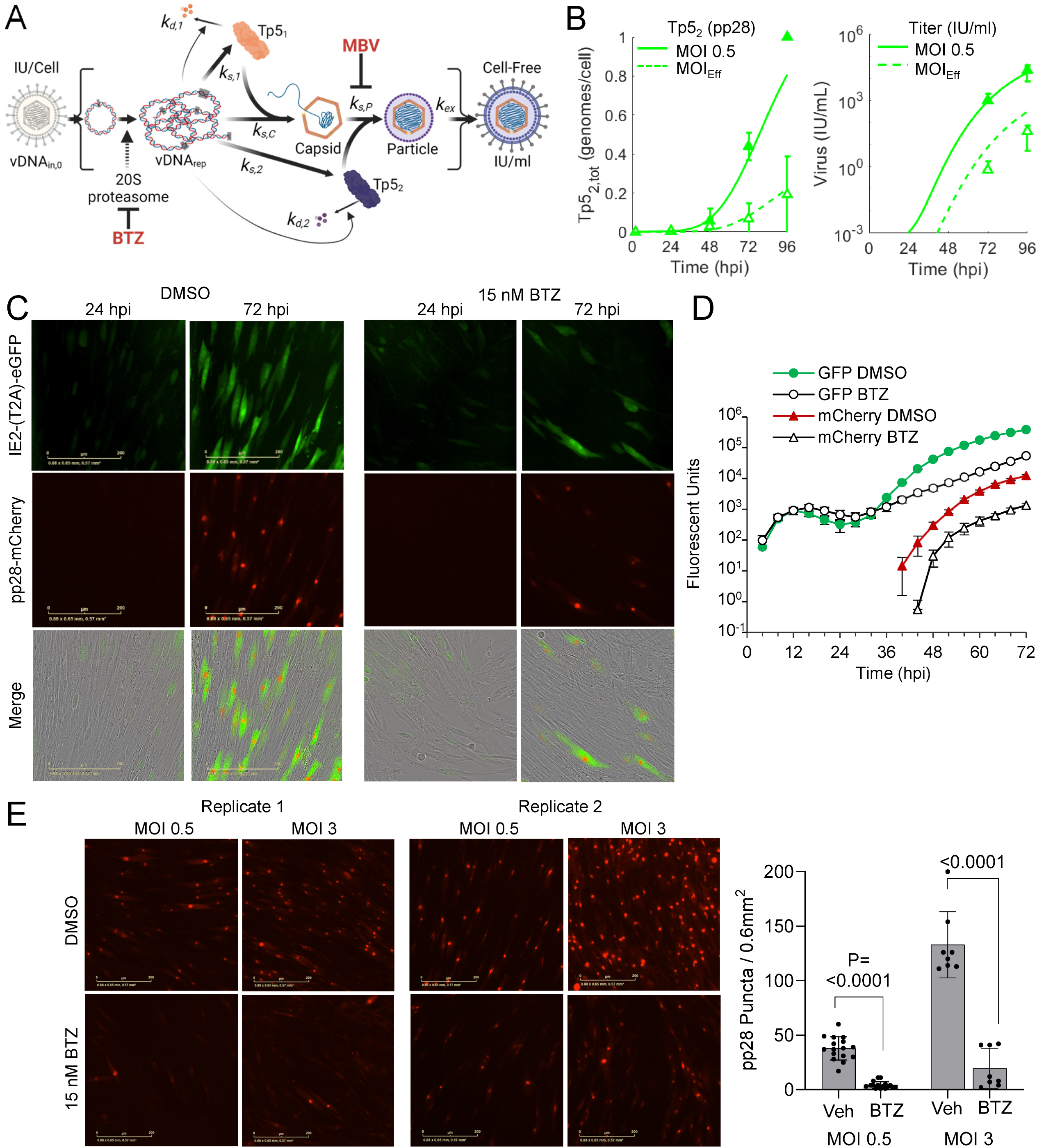
Live-cell imaging identifies timing of defect and reveals a subset of cells break inhibition, progressing to contain pp28-assembly compartments. **(A)** Schematic of a data-driven computational model of the late-lytic phase of HCMV replication indicating the location of bortezomib-mediated disruption between vDNA_in_ (input) and vDNA_rep_ (replicated). **(B)** Simulating bortezomib inhibition of infection at MOI 0.5 (solid line) by reducing vDNA_in_ (MOI 0.1; dashed line), and its impact on Tp5_2_ (pp28) levels and cell-free viral titers. Experimental data of pp28 levels (Fig. 2) and titers (Fig. 1) are overlayed for DMSO (closed triangles) and bortezomib-treated (open triangles) infections. Simulations suggest a possibility that proteasome inhibition reduces the total numbers of cells progressing to late lytic replication. **(C)** Live-cell imaging of infected fibroblasts. MRC-5 fibroblasts were grown to confluency and infected at multiple MOIs using recombinant HCMV TB40/E containing UL99-mCherry and IE2-(T2A)-eGFP which undergoes autocleavage producing free IE2 and eGFP. At 2 hpi, 15 nM bortezomib or DMSO vehicle was added and cultures were analyzed using IncuCyte Live-Cell Analysis System with images acquired every 4 hrs. Representative images shown from infections (0.5 IU/cell) at 24 and 72 hpi (see Fig. S1 avi files). **(D)** Integrated signal intensities per µm^2^ per image were measured over time for eGFP and mCherry during HCMV infection at 3.0 IU/cell, including standard error per image. **(E)** Distinct HCMV pp28-positive assembly compartments puncta were counted from each condition. Data are from 4-8 fields (0.6 mm^2^ per field) per indicated condition at 72 hpi with representative fields shown (n=2 biological replicates). Error bars represent standard deviation, and P values were calculated using a paired T test between the indicated samples.

To rule out this latter possibility, we expanded our experimental in vitro analyses using live cell imaging to provide a higher time resolution assessment and to observe single cell events. For these studies, we infected confluent fibroblasts using a BAC-derived HCMV recombinant expressing the IE gene, IE2 in-frame with eGFP and the L gene, UL99 (pp28) in-frame with mCherry (TB40e IE2-2A-eGFP UL99-mCh). The self-cleaving peptide motif, T2A was added between IE2 and eGFP reducing potential disruptions to IE2 activities and producing free eGFP. The use of this virus allows for monitoring the kinetics of IE (eGFP positive) and Late (mCherry positive) viral protein synthesis. Infections using TB40e IE2-2A-eGFP UL99-mCh virus were completed at MOIs of 0.5 and 3 IU/cell, and images were acquired every 4 hr for a 72 hr period. We added 15 nM bortezomib at 2 hpi without subsequent changes in culture media. As early as 4 hpi, we observed eGFP fluorescence in both vehicle and bortezomib-treated infections (**Fig. 7C**, **S1 videos**). However, by 72 hpi, proteasome inhibition resulted in fewer cells becoming eGFP-positive compared to vehicle. In cells with high eGFP signal, we also observed defined pp28-mCherry puncta which we interpret as cytoplasmic virus assembly compartments (**Fig. 7C**) (36). We quantified fluorescent signal intensities of eGFP and mCherry in the population overtime from 16 fields starting at 4 hpi for MOI 3, demonstrating delayed onset and reduced levels for both eGFP and mCherry in the cultures (**Fig. 7D**). To address per cell changes, we quantified pp28-mCherry puncta from multiple fields of view between vehicle and bortezomib conditions at both MOIs (**Fig. 7E**). These data show the presence of pp28-mCherry puncta under both conditions yet an approximate 87% reduction in the number of cells with puncta upon bortezomib treatment regardless of MOI (**Fig. 7E**). Overall, these data along with our simulations support a model in which bortezomib treatment reduced the MOI to MOI_Eff_ in which a subset of infected cells progress to contain cytoplasmic assembly compartments, and this is in contrast to the whole population uniformly progressing more slowly.

## DISCUSSION

Activities of proteasomes play an important role in the protracted and temporal stages required for lytic HCMV replication. Infection increases the levels of 20S subunits and peptidase activity starting after 24 hpi (8, 22). We have demonstrated the 20S proteasome reversible inhibitor, bortezomib disrupts viral replication. The addition of bortezomib at 2 hpi results in reduced levels of viral genomes by 24 hpi (**Fig. 1C**) and expression of proteins encoded by early and late viral genes (**Fig 2**). These results are consistent with studies by Tran et al. (8) using 2.5 µM MG132 which disrupts the 20S chymotrypsin-like and peptidyl glutamyl peptide hydrolyzing activities as well as calpain cysteine proteases (37). Those studies also demonstrated reductions in early and late HCMV transcripts occurs after 24 hpi. Because 20S proteasome activity likely contributes to multiple events, identifying precise mechanisms during HCMV replication are not possible. Our studies do support a previous conclusion that inhibiting the cellular proteasome has the greatest impact on viral DNA synthesis (22). In addition to quantifying a significant reduction in viral genomes (**Fig. 1C**), we observed reduced levels of proteins whose expression is influenced by genome copies with changes starting after 24 hpi (**Fig. 2C,D**). This includes pUL44 which functions as a viral DNA synthesis processivity factor. The conclusion is also supported by the timing of inhibitor addition as demonstrated in this work using bortezomib (**Fig. 4**) and studies using other inhibitors (8, 22). Addition of bortezomib starting at 48 hpi had limited effect on genome levels and virus production when compared to addition at 2 or 24 hpi (**Fig. 4B**). However, these observations do not exclude the possibility that the proteasome directly regulates viral gene expression as suggested by Tran et al. (8). Its role in regulating transcription is likely due to non-catalytic activities of the 19S regulator complex which is supported by studies from Winkler and Kalejta (17). They demonstrated depletion of 19S subunits results in a defect in RNA Pol II elongation at the HCMV major immediate early promoter (17).

The suppression of viral DNA synthesis upon proteasome inhibition is likely multifactorial. In addition to temporal changes in viral proteins, infection requires accompanied changes in cellular proteins and associated network processes. HCMV infection stimulates degradation restriction factors including Daxx, PML and BclAF1 (10–12, 38). These factors inhibit viral gene expression prior to 12 hpi, indirectly impacting viral DNA synthesis. However, infection upon knockdown of either PML or Daxx failed to abrogate the effects of MG132-mediated proteasome inhibition of synthesis suggesting the existence of other targets (22). We do expect that the viral protein-mediated regulation of cell cycle factors to be involved. Stimulation of infected cells into a G1/S-phase-like state contributes to the onset of viral DNA synthesis. Examples include HCMV pp71 which stimulates E2F activity by promoting proteasome-dependent degradation of Rb (10) and pUL21a which stimulates degradation of APC/C subunits and cyclin A (15, 16). HCMV IE2 also promotes an S-phase-like state (39–42), and we observed reduced IE2 expression by 48 hpi (**Fig. 2A,C**). Recent studies by Lin et al (43) highlight the impact of host protein turnover during HCMV infection using quantitative proteomics. These studies detected 64 proteins stabilized by 12 hpi during exposure with either MG123 or bortezomib, albeit at high concentrations, including host restriction factors and cell cycle regulators. These studies further support the hypothesis that manipulation of cellular proteosome is necessary at early times for efficient HCMV genome synthesis.

Bortezomib is an FDA approved proteasome inhibitor used in the treatment of multiple myeloma. Reactivation of latent herpesviruses VZV and, to a lesser extent, HCMV has been observed in multiple myeloma patients treated with bortezomib and, therefore, antiviral prophylaxis may be recommended (44, 45). In vitro, bortezomib treatment may also inhibit the lytic cycle of other herpesviruses. When treated with bortezomib, EBV-infected cells showed a reduced accumulation of viral DNA and proteins and produced less infectious virus (46). In HSV-1, bortezomib treatment may disrupt transport of the nucleocapsid to the nucleus and nucleolar remodeling (47). Our studies have shown that bortezomib also inhibits HCMV replication. In vitro, bortezomib treatment suppresses additional viruses including Dengue virus (48) and Chikungunya virus (49). These examples further highlight the critical role of host proteasomes in viral infection.

In virology, the use of MOI as a concentration of infectious unit (IU) or plaque forming unit (PFU) per cell to define an inoculum is standard practice. However, numerous limitations exist including, for example, when viral titers are determined on a different cells type than that used in experiments or variations in methods used to define an infectious particle between laboratories. Recently, we began using absolute quantification of viral genomes per cell (35); however, this too has its limitations as it is an average of the population. Based on the observations in these studies, we have introduced the term “effective MOI” or MOI_eff_ along with computational modeling to account for these differences (**Fig. 7**). We use the term to define the impact of proteasome inhibition on the initial input MOI as only a fraction of the cells progresses to fulminant infection. This MOI_eff_ has implication beyond these studies when it comes to selection and emergence of genetic variants. For HCMV, passage of epithelial/endothelial-tropic strains on different cell types selects for distinct populations (50–52), and passage of virus in the presence of ganciclovir and other anti-HCMV compounds selects for resistance. Inadequate dosing of ganciclovir has been attributed to emergence of antiviral resistance, and these changes have been documented under both in vitro and in vivo conditions (Reviewed in (53)). With bortezomib, it is conceivable that we can select unique variants showing resistance to proteasome inhibition and introducing new gain-of-function experimental tools to help uncover previously masked mechanisms.

Overall, our studies have demonstrated the impact of bortezomib on HCMV replication showing defects in viral DNA synthesis and downstream consequences when analyzing the population. We determined that bortezomib prevents replication-induced changes in cellular factors which likely synergizes with the HCMV kinase inhibitor, maribavir to block infection. Finally, we defined the reductions using real-time live cell imaging with *in-silico* analysis to be the result a subset of cells progressing to late stages of infection and leading to the introduction of the term, effective MOI.

## MATERIALS AND METHODS

### Cell culture and Viruses

MRC-5 fibroblasts (ATCC) were propagated in Dulbecco’s modified Eagle medium (DMEM) (Thermo Fisher Scientific) containing 7% fetal bovine serum (FBS) (Atlanta Biologicals) and 1% penicillin-streptomycin (Thermo Fisher Scientific). MRC-5s were grown to confluency at 37°C in 5% CO_2_ prior to infection. TB40/E GFP viral stocks were prepared by transfecting BAC DNA into MRC-5 fibroblasts with 1 µg of plasmid expressing UL82 (pCGN-pp71). Using a Gene Pulser Xcell electroporation system (Bio-Rad Laboratories) cells were electroporated in 4 mm gap cuvettes at 260 mV for 30 ms. Stocks of TB40e IE2-2A-eGFP UL99-mCh were prepared by infecting a monolayer of confluent MRC-5 cells at an MOI of 0.01. Cell culture medium was collected and then pelleted through a sorbitol cushion (20% sorbitol, 50 mM Tris-HCl, pH 7.2, 1 mM MgCl_2_) at 20,000xg for 90 min with a Sorvall WX-90 ultracentrifuge and SureSpin 630 rotor (Thermo Fisher Scientific). Titers of viral stocks were determined with MRC-5 cells using a limiting dilution assay (50% tissue culture infectious dose [TCID50]) in 96-well dishes. Cells were counted prior to infection using a hemocytometer and infected using a multiplicity of infection (MOI) of 0.5 IU/cell unless otherwise noted. At 2 hpi cells were washed with phosphate-buffered saline (PBS) and the media was replaced. Drug treatments were performed at 2 hpi and drug were changed every 24 hours unless otherwise noted. Cells were treated with bortezomib (BTZ) (Sigma Aldrich), maribavir (MBV)(Takeda, formerly Shire), or DMSO as a vehicle control. Titers of the resulting culture media were analyzed by plating serial dilutions of culture media for 2 hpi, replacing with fresh media at 2 hpi, staining for IE1 at 48 hpi, and counting IE1-postive cells per well at a given volume plated resulting in infectious units (IU) per ml.

Fluorescently tagged TB40/E expressing IE2-2A-eGFP and UL99-mCh was generated from HCMV BAC-derived clinical strain TB40/E. The virus was cloned using *galK* BAC recombineering methods, described previously (54). For the IE2-2A-eGFP modification *galK* was inserted at the C terminus of UL122 with the following primers: forward, 5′-<u>T</u>GAGCCTGGCCATCGAGGCAGCCATCCAGGACCTGAGGAACAAGTCTCAGCCTGTTGACAATTAATCATCGGCA-3′, and reverse, 5′-CACGGGGAATCACTATGTACAAGAGTCCATGTCTCTTTCCAGTTTTTCACTCAGCACTGTCCTGCTCCTT-3′. Note that this insertion deletes the stop codon of UL122. *GalK*+ recombinants were selected on plates in which galactose was the only carbon source. Positive recombination was validated by PCR. A second recombination was performed to remove the *GalK* cassette and replace it with a cassette containing the T2A element, a “self-cleaving” peptide sequence derived from Thosea asigna virus (55), with the sequence 5′-GGTTCAGGTGAGGGCAGAGGCTCACTCTTGACGTGCGGTGATGTAGAAGAAAACCCCGGTCCT-3′, which was cloned in-frame with eGFP and amplified with the following primers: forward, 5′-TGAGCCTGGCCATCGAGGCAGCCATCCAGGACCTGAGGAACAAGTCTCAGGGTTCAGGTGAGGGCAGAGGCTCA–3′, and reverse, 5′- CACGGGGAATCACTATGTACAAGAGTCCATGTCTCTTTCCAGTTTTTCACTTACTTGTACAGCTCGTCCATGCC-3′, containing sequence homology to the 3′ end of UL122. Purified PCR product was transformed into electrocompetent SW105 cells. Clones were selected for loss of *galK* expression on plates containing 2-deoxy-D-galactose, then screened by PCR analysis and sequenced to verify the position and integrity of the inserted cassette. In a similar manner, *galK* was inserted into the UL99 locus using primers: forward, 5’-CAACGTCCACCCACCCCCGGGACAAAAAAGCCCGCCGCCCCCTTGTCCTTTCCTGTTGACAATTAATCATCGGCA-3’ and reverse, 5’- GTGTCCCATTCCCGACTCGCGAATCGTACGCGAGACCTGAAAGTTTATGAGTCAGCACTGTCCTGCTCCTT-3’. *GalK*-positive clones were grown and reverted with a PCR product generated from the following primers forward, 5’- CAACGTCCACCCACCCCCGGGACAAAAAAGCCCGCCGCCCCCTTGTCCTTTGTGAGCAAGGGCGAGGAGCTGTTCACCG-3’ and reverse, 5’- GTGTCCCATTCCCGACTCGCGAATCGTACGCGAGACCTGAAAGTTTATGAGTTACTTGTACAGCTCGTCCATGCCGAGAGT-3’ using mCherry as a template. Recombinant clones were counter selected, screened by PCR, and sent for confirmatory sequence analysis.

### Incucyte Live Cell Imaging

MRC-5 fibroblasts were grown to confluency and infected at 0.5 IU/cell or 3.0 IU/cell using recombinant HCMV TB40/E containing IE2-(T2A)-eGFP and UL99-mCherry. At 2 hpi, 15 nM bortezomib or DMSO vehicle was added, and wells were analyzed using IncuCyte Live-Cell Analysis System (Sartorius) at 37°C in 5% CO_2_. For each condition and replicate, 16 images including brightfield, green, and red channels were taken every 4 hrs at 20x magnification to 72 hpi. Movie Avi files are provided in supplemental materials.

### Nucleic Acid and Protein Analyses

Protein levels were analyzed via western blotting. Cells were resuspended in Protease Inhibitor Cocktail (Sigma), HALT Phosphatase Inhibitor Cocktail (Thermo Fisher Scientific), and lysis buffer (1.5% SDS, 10 mM NaCl, 50 mM Tris Cl (pH 7.2), 1 mM EDTA) and lysed by sonication. Total protein quantity in samples was determined via Pierce BCA Protein Assay Kit (Thermo Fisher Scientific). Laemmli sample buffer was added to samples. Protein samples were resolved by stain-free sodium dodecyl sulfate–10% polyacrylamide gel electrophoresis (SDS-PAGE) and transferred to a nitrocellulose membrane (GE Healthcare) using the semi-dry Trans-Blot turbo transfer system (Bio-Rad). The membrane was blocked in 5% milk in TBST (TBS, 0.1% Tween 20) for 1 hour at room temperature, incubated in primary antibody overnight at 4°C with 5% milk or 5% bovine serum albumen in TBST, then incubated for 1 hour with goat anti mouse IgG conjugated to horseradish peroxidase or 2 hours with donkey anti rabbit IgG conjugated to horseradish peroxidase at room temperature in 5% milk in TBST. HRP-conjugated antibodies were detected using enhanced chemiluminescence reagents (ECL) (Thermo Fisher Scientific) and ChemiDoc MP Imaging System (BioRad).

The following antibodies were used for western blot at the dilution indicated: mouse anti-p21 (clone CP36, 1:1000, Millipore-Sigma), rabbit anti-Cyclin B1 (clone D5C10, 1:1000, Cell Signaling Technology), mouse anti-Cdk1 (clone POH1, 1:500, Cell Signaling Technology), mouse anti-PCNA (clone PC10, 1:1000, Cell Signaling Technology), mouse anti-Lamin A/C (clone 4C11, 1:2000, Cell Signaling Technology), rabbit anti-phospho-Lamin A/C (Ser22) (1:1000, Cell Signaling Technology), mouse anti pUL44 (clone 10D8, 1:10,000, Virusys). Secondary antibodies were Peroxidase AffiniPure Donkey Anti-Rabbit IgG (1:10,000, Jackson Immuno Research) and Peroxidase AffiniPure Goat Anti-Mouse IgG (1:10,000, Jackson Immuno Research). The HCMV antibodies mouse anti-IE1 (clone 1B12, 1:1000), mouse anti-IE2 (clone 3A9, 1:1000), mouse anti-pp28 (clone 10B4-29, 1:1000) were generously provided by Tom Shenk (Princeton University).

Cellular and viral DNA and RNA contents were determined using quantitative PCR (qPCR) analysis. Cells were collected, resuspended in lysis buffer (400 mM NaCl, 10 mM Tris, pH 8.0, 10 mM EDTA, 0.1 mg/ml proteinase K, 0.2% SDS), and incubated at 37°C overnight. DNA was extracted using phenol-chloroform and precipitated using ethanol. Alternatively, cells were collected, and DNA extracted using the DNeasy Blood & Tissue Kit (Qiagen). qPCR was completed using primers for HCMV UL123, Gene ID 3077513 (5′- GCCTTCCCTAAGACCACCAAT-3′ and 5′-ATTTTCTGGGCATAAGCCATAATC-3′) and cellular TUBB, Gene ID: 203068, (5’-GCAGAAGAGGAGGAGGATTTC-3’ and 5’-CAGTTGAGTAAGACGGCTAAGG-3’). Quantification was performed using Power SYBR Green PCR Master Mix (Thermo Fisher Scientific) and QuantStudio 6 Flex real-time PCR System (Thermo Fisher Scientific). An arbitrary standard curve was used to determine relative quantities of DNA. Within an experiment each primer set had an arbitrary standard curve and was normalized to TUBB. For RNA studies, total RNA was isolated using the RNeasy Mini Kit (Qiagen). 1 µg RNA was treated with the TURBO DNA-free Kit (Thermo Fisher Scientific) and used to synthesize cDNA with random hexamers (Thermo Fisher Scientific) and Superscript III reverse transcriptase (Thermo Fisher Scientific). qPCR was performed as described using primers against P21, Gene ID: 1026 (5’-TTAGCAGCGGAACAAGGAGTCAGA-3’ and 5’- ACACTAAGCACTTCAGTGCCTCCA-3’) and GAPDH, Gene ID: 2597 (5’- CTGTTGCTGTAGCCAAATTCGT-3’ and 5’-ACCCACTCCTCCACCTTTGAC-3’). An arbitrary standard curve was used to determine relative quantities of RNA. Within an experiment each primer set had an arbitrary standard curve and was normalized to GAPDH.

### In silico simulations

For simulating the effects of BTZ on HCMV replication, we utilized the recently developed model of Monti *et al.* (35). Briefly, this model uses a linear relationship between MOI and input vDNA at 2 hpi (vDNA_in,0_) as input for an empiric function to simulate vDNA over time, which then drives a mechanistic, ordinary differential equation (ODE)-based model of late HCMV replication. Given MOI 0.5 and 3 used in this manuscript, we converted these values to vDNA_in,0_ using the linear relationship (35) and then simulated predicted protein and viral titer levels and compared these results to experimental data. Notably, the viral titer simulations were vertically scaled by 5 x 10^4^ IU/ml to account for the unit differences between our normalized model and viral titer data. For calculating effective MOI (*MOI_eff_*,), the number of infected cells is related to MOI via the Poisson distribution (56) where the number of cells infected with one or more viruses is:

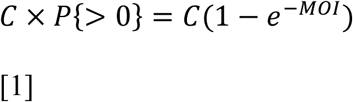

Where *C* is the number of cells and *MOI* is the intended MOI of the infection. Given the observed ∼80% reduction in number of infected cells in the presence of BTZ, an effective MOI (*MOI_eff_*) can be generated:

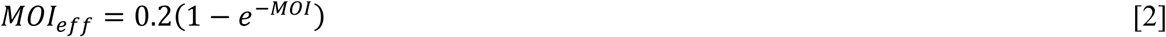

From *MOI_eff_*, vDNA_in,0_ can be calculated and then input into the model and compared to data. All model simulations were performed in MATLAB (MathWorks) using code available at https://github.com/MCWComputationalBiologyLab/Monti_2022_PNAS.

### Drug Synergism and Statistic Analyses

Drug synergism combination index (CI) values were determined using the CompuSyn software tool (57). Experiments were analyzed using paired Student’s t-test or ANOVA with multiple comparisons with GraphPad Prism software. P < 0.05 was considered significant.

## Data Availability

All data and reagents from these studies will be made available upon request.

## ACKNOWLEDGEMENTS

We thank Dr. Tom Shenk for providing antibodies against HCMV proteins. Also, we thank Terhune laboratory members and the labs of Drs. Amy Hudson and Ravit Boger for their discussion and feedback on these studies. Research reported in this publication was supported by the National Institute of Allergy and Infectious Diseases division of the National Institutes of Health under award number R01AI083281 to S.S.T., award number R21AI149039 to S.S.T. and R.K.D., award number F30AI179084 to C.E.M., and award numbers R01AI155979 and R21AI147152 to E.A.M. These studies have also been supported by a generous philanthropic gift by The Stead Family Foundation. The funders had no role in study design, data collection and interpretation, or the decision to submit the work for publication.

Author contributions were as follows: K.M.C. and S.S.T., conceptualization and methodology; K.M.C. with assistance from K.L.R., investigation; C.E.M. and R.K.D., mathematical modeling; S.S.T., R.K.D., and E.A.M, funding acquisition and supervision; K.M.C. and S.S.T., formal analysis; K.M.C., C.E.M, and S.S.T., visualization; K.M.C. and S.S.T., writing - original draft; K.M.C., C.E.M., R.K.D., E.A.M., and S.S.T., writing - review & editing. All authors have read and approved the final manuscript.

